# Function and clinical relevance of RHAMM isoforms in pancreatic tumor progression

**DOI:** 10.1101/598334

**Authors:** Soyoung Choi, Dunrui Wang, Xiang Chen, Laura H. Tang, Akanksha Verma, Zhengming Chen, Bu Jung Kim, Leigh Selesner, Kenneth Robzyk, George Zhang, Sharon Pang, Teng Han, Chang S. Chan, Thomas J. Fahey, Olivier Elemento, Yi-Chieh Nancy Du

## Abstract

The receptor for hyaluronic acid-mediated motility (RHAMM) is upregulated in various cancers. We previously screened genes upregulated in human hepatocellular carcinomas for their metastatic function in a mouse model of pancreatic neuroendocrine tumor (PNET) and identified that human *RHAMM*^*B*^ promoted liver metastasis. It was unknown whether *RHAMM*^*B*^ is upregulated in pancreatic cancer or contributes to its progression. In this study, we found that RHAMM protein was frequently upregulated in human PNETs. We investigated alternative splicing isoforms, *RHAMM*^*A*^ and *RHAMM*^*B*^, by RNA-Seq analysis of primary PNETs and liver metastases. *RHAMM*^*B*^, but not *RHAMM*^*A*^, was significantly upregulated in liver metastases. RHAMM^B^ was crucial for *in vivo* metastatic capacity of mouse and human PNETs. RHAMM^A^, carrying an extra 15-amino acid-stretch, did not promote metastasis in spontaneous and experimental metastasis mouse models. Moreover, *RHAMM*^*B*^ was substantially higher than *RHAMM*^*A*^ in pancreatic ductal adenocarcinoma (PDAC). *RHAMM*^*B*^, but not *RHAMM*^*A*^, correlated with both higher *EGFR* expression and poorer survival of PDAC patients. Knockdown of EGFR abolished RHAMM^B^-driven PNET metastasis. Altogether, our findings suggest a clinically relevant function of *RHAMM*^*B*^, but not *RHAMM*^*A*^, in promoting PNET metastasis in part through EGFR signaling. *RHAMM*^*B*^ can thus serve as a prognostic factor for pancreatic cancer.

## Main text

Metastasis accounts for 90 percent of cancer deaths. We developed a mouse model of well-defined multistage tumorigenesis: *RIP-Tag; RIP-tva* to identify metastatic factors [1]. We identified that the receptor for hyaluronic acid (HA)-mediated motility, isoform B (RHAMM^B^), significantly promotes liver metastasis of pancreatic neuroendocrine tumors (PNET) in *RIP-Tag; RIP-tva* mouse models [2]. Expression of RHAMM is restricted in normal adult tissues, but is upregulated in cancers [3, 4]. Increased production of glycosaminoglycan, HA, is correlated with increased migration and invasion in aggressive cancers [5]. CD44 and RHAMM are two major HA receptors. The roles of CD44 isoforms in cancer have been studied extensively, but the functions of RHAMM isoforms in tumorigenesis are less clear. *RHAMM* encodes 18 exons and alternative splicing generates different isoforms. *RHAMM*^*A*^ includes all exons and *RHAMM*^*B*^ lacks exon 4 (Fig. 1A). Here we aimed to determine the clinical relevance of *RHAMM*^*A*^ and *RHAMM*^*B*^ isoforms and their functions in pancreatic cancer.

**Fig. 1.**
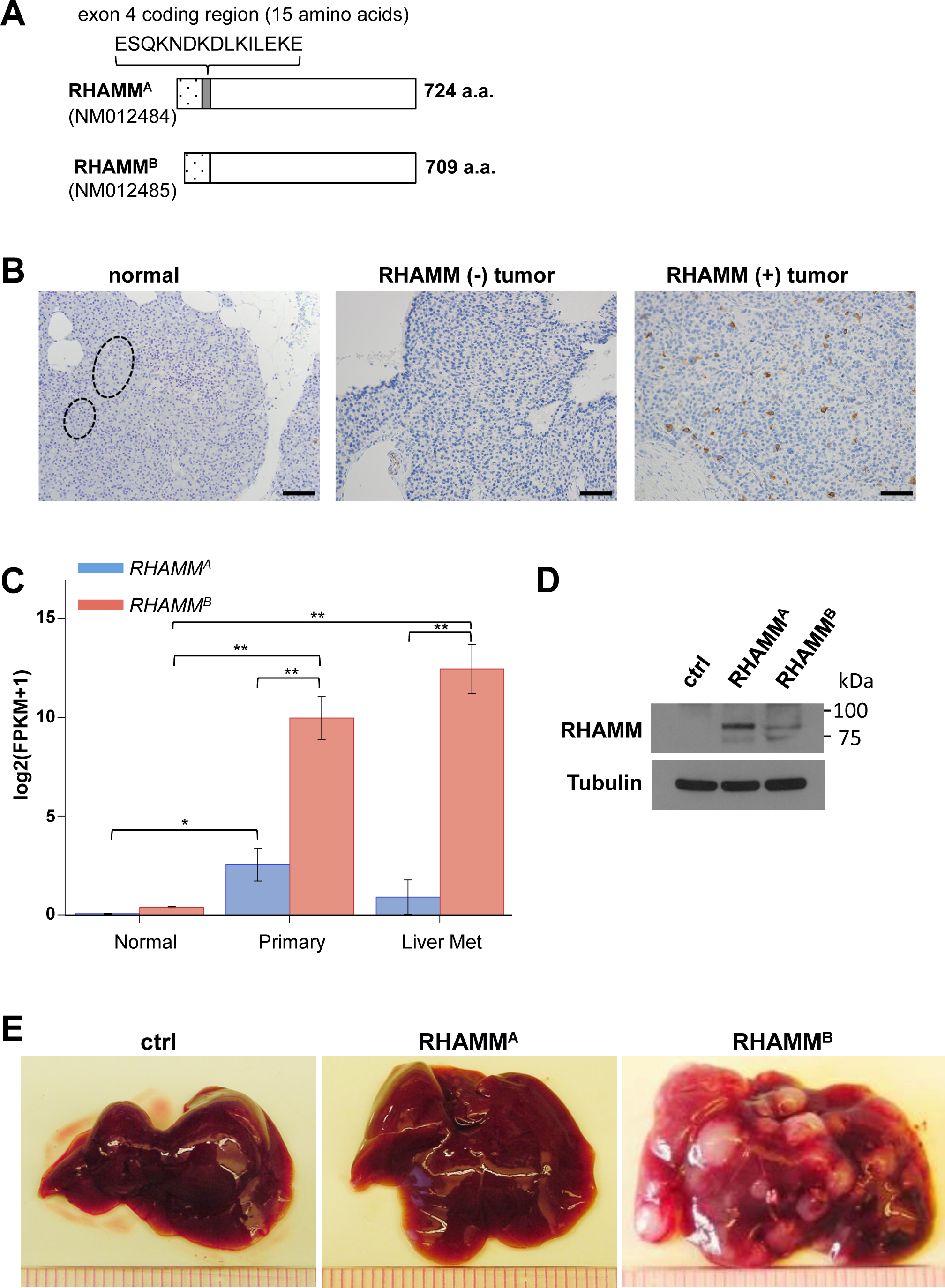
RHAMM^B^, but not RHAMM^A^, is upregulated in human PNETs and promotes liver metastasis of mouse PNET cells. (A) Diagram of RHAMM^A^ and RHAMM^B^ proteins. (B) RHAMM is upregulated in 54 of 83 cases (65%) of human PNETs in immunohistochemical staining. Left: Normal pancreas with islets in dashed circle. Middle: RHAMM negative PNET. Right: RHAMM positive PNET. Original magnification: 20X. Scale bar, 50 µm. (C)RNA-seq analysis showed that *RHAMM*^*B*^ is significantly upregulated compared to *RHAMM*^*A*^ in primary human PNETs and liver metastases. The p value was calculated using two-way ANOVA followed by Tukey’s test. **: p < 0.0001, *: p < 0.05. Error bars represent standard error of mean. (D) Western blot analysis of human RHAMM in mouse N134 cell line (control), N134-RHAMM^A^ cells, and N134-RHAMM^B^ cells. (E) A total of 1 million N134 cells, N134-RHAMM^A^ cells, or N134-RHAMM^B^ cells were injected into the tail vein of NSG mice (n = 5 for each group). Five weeks later, the recipient mice were euthanized to survey for metastatic sites and incidence. Representative liver photos were shown.

### RHAMM^B^, but not RHAMM^A^, is upregulated in human PNET liver metastases

To investigate RHAMM expression in human PNETs, a tissue microarray consisting of 83 PNETs was immunostained for RHAMM. RHAMM was not detectable in the normal pancreas, while 54 of 83 (65%) PNETs exhibited cytoplasmic staining using an antibody that recognizes common region in RHAMM isoforms (Fig. 1B). Because isoform-specific RHAMM antibodies were not available, we investigated the mRNA levels of *RHAMM*^*A*^ and *RHAMM*^*B*^ by performing RNA-Seq analysis on 27 primary PNETs and 12 liver metastases, using 89 human islets from Gene Expression Omnibus database for comparison. Consistent with our immunohistochemical data, normal islets had very low *RHAMM*^*A*^ and *RHAMM*^*B*^ mRNA (Fig. 1C). *RHAMM*^*B*^ was significantly higher than *RHAMM*^*A*^ in both primary and metastatic PNETs, suggesting that *RHAMM*^*B*^ was the predominant isoform naturally expressed in PNETs. Although *RHAMM*^*A*^ levels in primary tumors were significantly higher than those in normal islets (p = 0.0002), *RHAMM*^*A*^ levels in metastatic tumors were not significantly higher than those in normal islets (p = 0.8928). The mRNAs of *RHAMM*^*A*^ and *RHAMM*^*B*^ were readily detectable in additional primary PNETs and metastases by RT-qPCR using isoform-specific primers (Fig. S1A-B).

We compared metastatic potential of RHAMM^A^ to RHAMM^B^ in *RIP-Tag; RIP-tva* models of spontaneous metastasis and tail vein assays [1, 2]. In contrast to RHAMM^B^, RHAMM^A^ did not promote spontaneous metastasis (Table S1). Then, we generated N134 cells overexpressing RHAMM^A^ (N134-RHAMM^A^). N134 is a cell line derived from a PNET of *RIP-Tag; RIP-tva* mouse [1]. Although there were more RHAMM^A^ than RHAMM^B^ for unknown reasons (Fig. 1D), only one visible tumor was found in 5 immunodeficient NOD/scid-IL2Rgc knockout (NSG) mice receiving N134-RHAMM^A^ cells after 5 weeks, while all 5 mice receiving N134-RHAMM^B^ cells developed large liver metastases within 5 weeks (Fig. 1E and Fig. S2A). To detect micrometastases, we performed immunostaining for synaptophysin, a neuroendocrine marker. Mice receiving N134 cells and N134-RHAMM^A^ cells had an average of 1.8 and 0.6 liver micrometastases, respectively (Fig. S2B). These data suggest that the unique 15-amino acid-stretch, ESQKNDKDLKILEKE, which is present in RHAMM^A^ but not in RHAMM^B^, inhibited the metastatic function of RHAMM.

### RHAMM^B^ is crucial for the metastatic potential of human PNET cell line BON1-TGL

BON1 is the most utilized human PNET cell line and was established from a peri-pancreatic lymph node in a patient with metastatic PNET. We found that BON1 had much higher expression of *RHAMM*^*B*^ than *RHAMM*^*A*^ as determined by RNA-Seq (Fig. 2A). We performed shRNA-mediated knockdown of total *RHAMM* to investigate whether this reduces metastasis of BON1-TGL cells, which carry the thymidine kinase/*green fluorescent protein /luciferase* fusion reporter (TGL). Knockdown of *RHAMM* by shRNA was confirmed (Fig. 2B-C). We used an orthotopic model of PNET liver metastases by injecting cells into the spleen of NSG mice [6]. Mice receiving control cells developed an average of 82.5 liver metastases after 3 weeks, while mice receiving BON1-TGL-*shRHAMM* cells developed an average of 17 liver metastases with significantly lowered tumor burden (Fig. 2D-F).

**Fig. 2.**
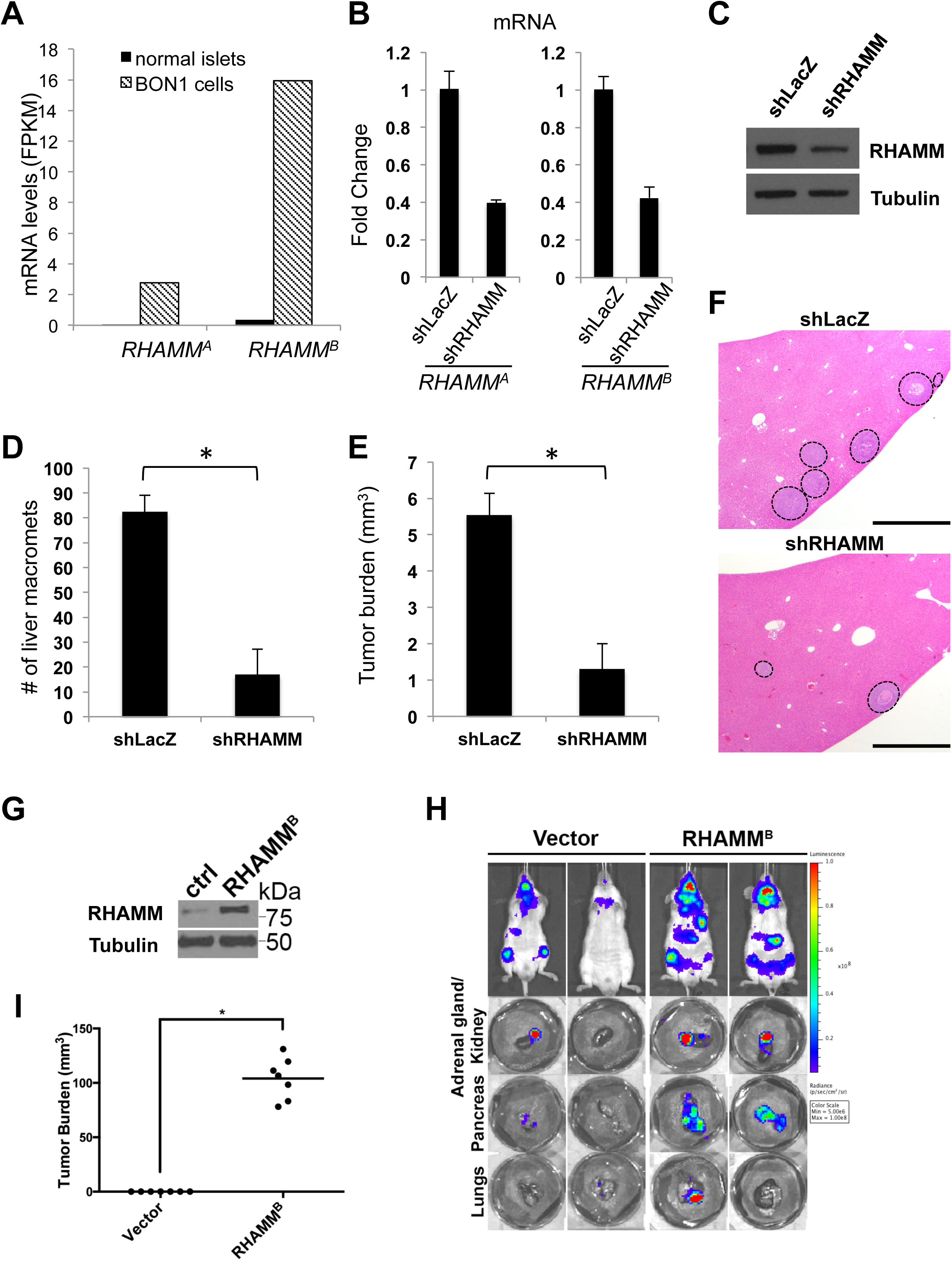
RHAMM^B^ is crucial for metastatic potential of human PNET cell line, BON1-TGL. (A) *RHAMM*^*A*^ and *RHAMM*^*B*^ expression in BON1 cell line compared to those in normal islets. (B) *RHAMM*^*A*^ and *RHAMM*^*B*^ knockdown in BON1-TGL cell line by sh*RHAMM* as determined by RT-qPCR analysis. (C) Western blot analysis of RHAMM and tubulin (as a loading control) in BON1-TGL-*shLacZ* cell line and BON1-TGL-*shRHAMM* cell line. (D-F) RHAMM knockdown greatly inhibited liver metastasis of BON1-TGL cells. A total of 0.5 million each BON1-TGL-*shLacZ* (control) or BON1-TGL-*shRHAMM* were injected into the spleen of NSG mice (n = 4 for each group). After 3 weeks, the recipient mice were euthanized to survey for metastatic sites and incidence. The number of liver macrometastases (D) and the tumor burden of liver macrometastases (E) were recorded. *: statistically significantly different (p < 0.05, one-tailed Mann-Whitney U test). Error bars in this figure represent standard deviation. (F) Liver sections with hematoxylin and eosin stain. Dashed circles indicate metastases. Original magnification: 10X. Scale bar, 1 mm. (G) RHAMM^B^ overexpression in BON1-TGL-RHAMM^B^ cell line. Western blot analysis of RHAMM and tubulin (as a loading control) are shown. (H) Representative bioluminescent images of NSG mice 4 weeks after injection (upper panel) and their organs (lower panel). A total of 1 million cells was injected into NSG mice via intracardiac injection (n = 7). (I) The tumor burden of macrometastases at adrenal glands was documented. *: statistically significantly different, p<0.05, t-test.

To determine whether even higher RHAMM^B^ levels would further enhance metastasis of BON1-TGL cells, we generated BON1-TGL cells overexpressing RHAMM^B^ (Fig. 2G). We injected cells into NSG mice via intracardiac injection. The increased levels of RHAMM^B^ enhanced metastasis of BON1-TGL in mice throughout the mouse body as visualized by bioluminescence imaging and signals from multiple organs were higher in mice receiving BON1-TGL-RHAMM^B^ than those in mice receiving BON1-TGL overexpressing a control vector (Fig. 2H). Notably, we observed macrometastases at the adrenal glands of mice receiving BON1-TGL-RHAMM^B^, but not in control mice (Fig. 2I). Taken together, RHAMM^B^ is crucial for PNET metastasis.

### *RHAMM*^*B*^ is upregulated in human pancreatic ductal adenocarcinoma (PDAC) and correlates with poor survival

PDAC is the most common pancreatic cancer type. It was shown that total *RHAMM* is upregulated in primary PDAC by RT-qPCR of 14 matched tumors and adjacent normal tissues [7]. To compare *RHAMM*^*A*^ and *RHAMM*^*B*^ levels in PDAC, we analyzed publicly available The Cancer Genome Atlas (TCGA) datasets. Both *RHAMM*^*A*^ and *RHAMM*^*B*^ were expressed at significantly higher levels in PDAC than in normal pancreatic tissues, and *RHAMM*^*B*^ was substantially higher than *RHAMM*^*A*^ (Fig. 3A). Survival analysis showed that high *RHAMM* levels were correlated with a worse outcome (Fig. 3B). Furthermore, patients with high *RHAMM*^*B*^ had inferior survival compared to those with high *RHAMM*^*A*^ (Fig. 3C-D), suggesting that *RHAMM*^*B*^, but not *RHAMM*^*A*^, is a prognostic factor for survival of PDAC patients. Due to limited information available, we could not perform survival analysis for PNETs.

**Fig. 3.**
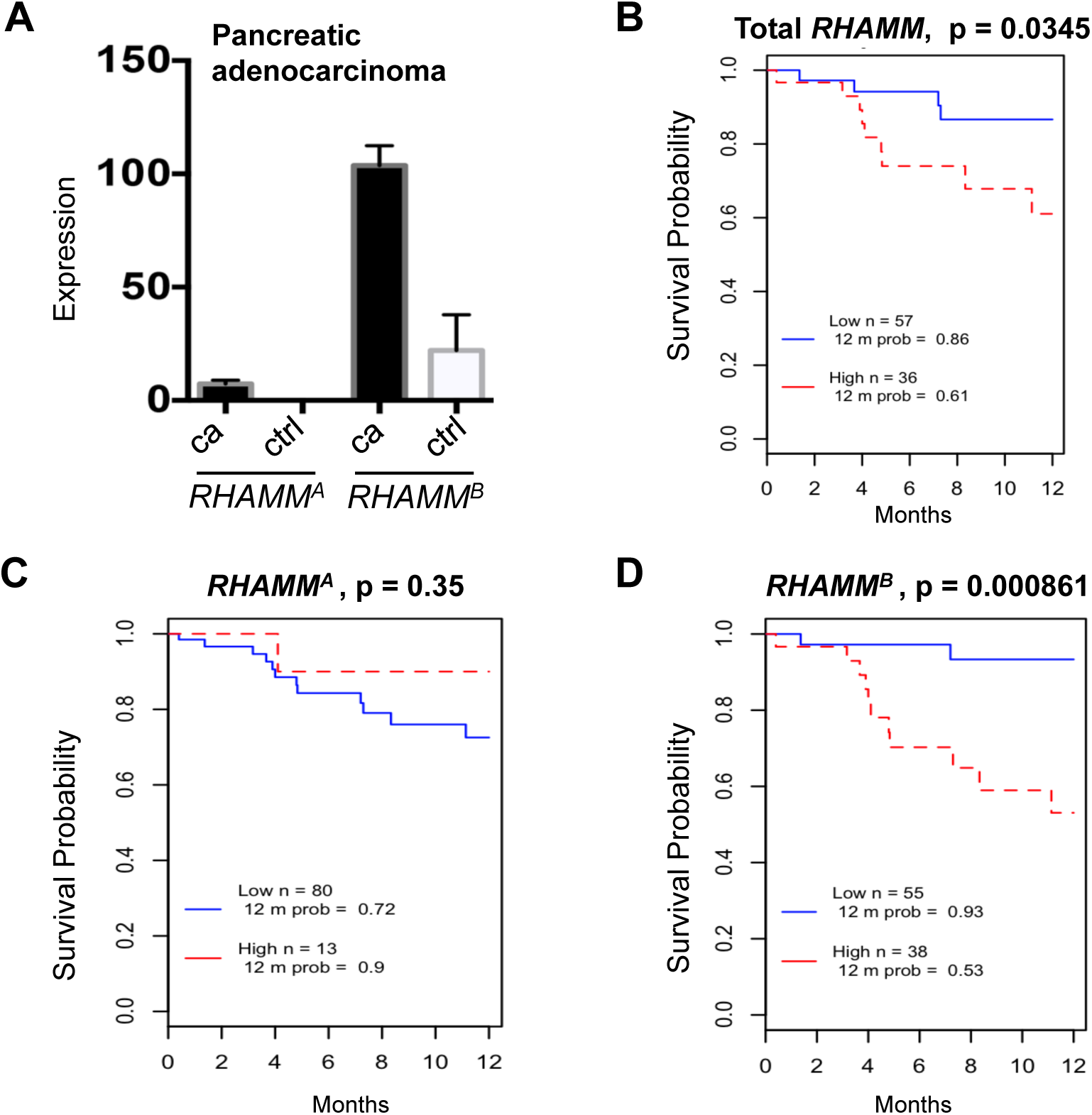
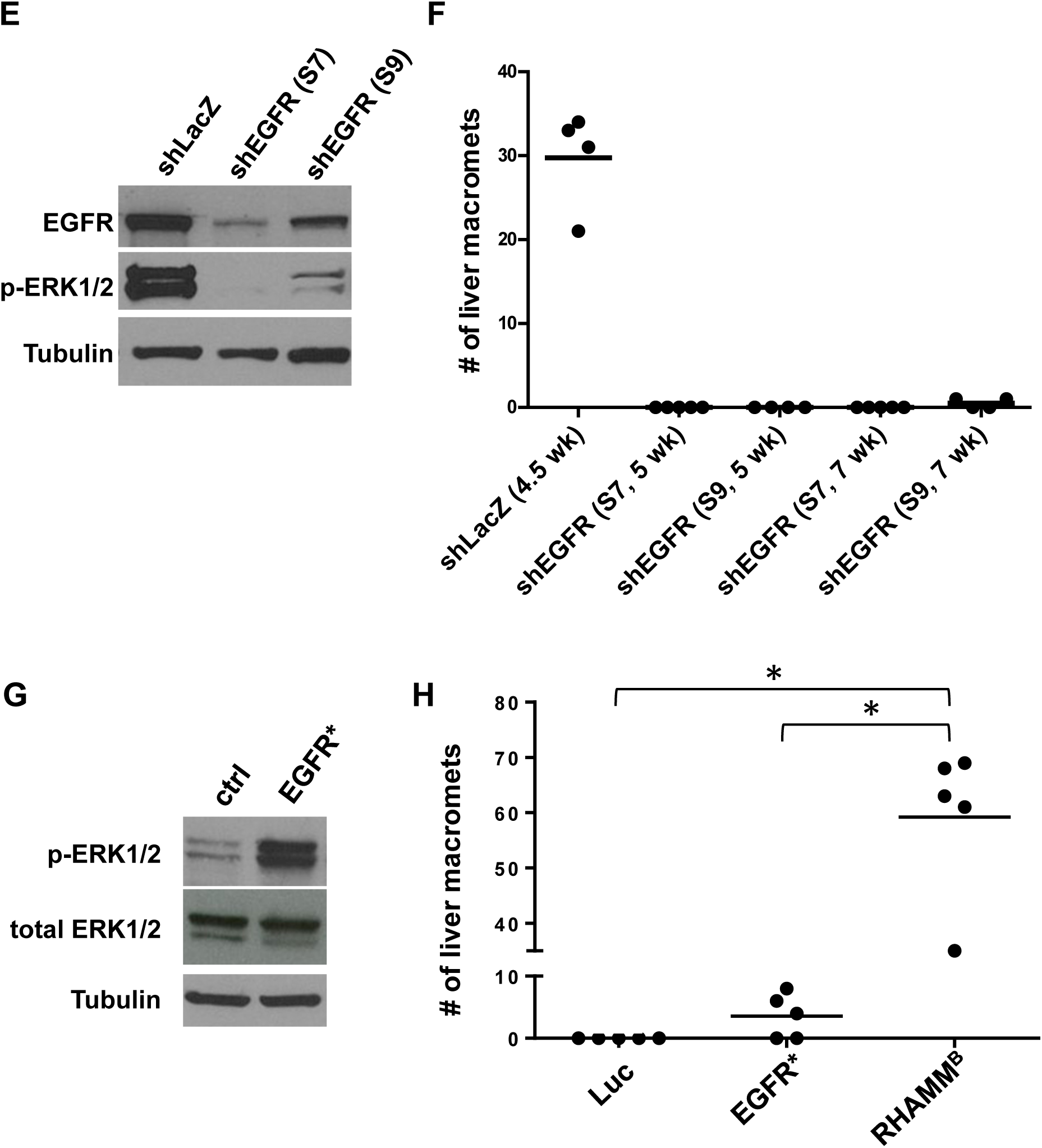
*RHAMM*^*B*^ is upregulated in human pancreatic ductal adenocarcinoma (PDAC) and correlates with poor survival. (A) *RHAMM*^*A*^ and *RHAMM*^*B*^ expression values from TCGA PDAC dataset. *RHAMM*^*A*^: uc003lzf or NM_012484. *RHAMM*^*B*^: uc003lzg or NM_012485. cancer (ca): n = 124; control (ctrl): n = 4. Bars and error bars represent means and standard errors. (B-D) Kaplan-Meier survival analysis of TCGA cohort with 93 PDAC cases. High and low represent the status of the *RHAMM* mRNA expression levels compared to average values. (E) Knockdown efficiency of two EGFR shRNAs. S7 and S9 reduced EGFR protein expression and p-Erk1/2 levels in N134 cells overexpressing RHAMM^B^ by Western blot analysis. α-tubulin was used a loading control. (F) EGFR knockdown greatly inhibited the liver metastasis of N134 cells overexpressing RHAMM^B^. A total of 1 million N134_RHAMM^B^_shLacZ, N134_RHAMM^B^_*shEGFR*(S7), or N134_RHAMM^B^_*shEGFR*(S9) cells were injected into the tail vein of NSG mice (n = 5 for each group). At the indicated time points, the recipient mice were euthanized to survey for metastatic sites and incidence. The number of liver macrometastases was recorded. (G) Western blot analysis of p-Erk1/2 and total Erk from N134 overexpressing luciferase (control), and N134_EGFR*. (H) N134 cells overexpressing luciferase (Luc), N134_EGFR* (EGFR*), N134_RHAMM^B^ (RHAMM^B^) were injected into the tail vein of NSG mice (n = 5, each group). Five weeks later (for Luc and EGFR* groups) or when mice were lethargic (for RHAMM^B^), mice were euthanized to survey for metastatic sites and incidence. *: p< 0.0001, One-way ANOVA and pairwise comparison with Tukey’s adjustment.

### EGFR signaling is required for RHAMM^B^-induced metastasis

EGFR activation is associated with worse survival in many malignancies. Analysis of TCGA PDAC dataset showed a good correlation between *EGFR* and *RHAMM*^*B*^ expression, but not between *EGFR* and *RHAMM*^*A*^ expression (Fig. S3A-B). We previously showed that EGFR signaling is activated in N134-RHAMM^B^ cells and an EGFR inhibitor, gefitinib, induces apoptosis of N134-RHAMM^B^ cells [2]. These findings led us to hypothesize that enhanced EGFR signaling involves RHAMM^B^-induced metastasis. To test this hypothesis, we used two different shRNAs targeting mouse *EGFR* (S7 and S9) as well as a control shRNA targeting LacZ (*shLacZ*). sh*EGFR*(S7 and S9) decreased the levels of both EGFR and p-ERK1/2, a downstream target of EGFR signaling, with S7 exhibiting a better knockdown efficiency than S9 (Fig. 3E).

In tail vein metastasis assays, mice receiving N134-RHAMM^B^-*shLacZ* became lethargic, showing a mean of 29.8 liver macrometastases per mouse within 5 weeks, but mice injected with N134-RHAMM^B^-*shEGFR*(S7) and N134-RHAMM^B^-*shEGFR*(S9) cells did not develop metastases 5 weeks post-injection (Fig. 3F). An additional cohort of mice injected with N134-RHAMM^B^-*shEGFR*(S7) still did not develop metastases 7 weeks post-injection, and only a mean of 0.5 liver macrometastases appeared in mice injected with N134-RHAMM^B^-*shEGFR*(S9) at this time point (Fig. 3F).

We investigated whether a constitutively active form of EGFR, EGFR*, is sufficient to recapitulate RHAMM^B^ activity in metastasis. Five of the 8 *RIP-Tag; RIP-tva* mice receiving RCASBP-*EGFR\** developed pancreatic lymph node metastases (62.5%) and 2 developed liver metastases (25%) (Table S1). We generated N134 cells overexpressing EGFR* (N134-EGFR*) for experimental metastasis (Fig. 3G). While no metastasis was found in mice receiving control N134-Luciferase after 6 weeks, an average of 6 liver macrometastases was detected in mice receiving N134-EGFR* (Fig. 3H). Although EGFR* promotes liver metastasis of PNETs in both spontaneous and experimental metastasis mouse models, the degree of metastasis is less than that of RHAMM^B^ (Table S1 and Fig. 3H). Therefore, these EGFR knockdown and EGFR* overexpression data suggest that activation of EGFR contributes to RHAMM^B^-induced PNET metastasis, but EGFR* cannot fully recapitulate the metastatic phenotype of RHAMM^B^. Further studies are required to identify signals other than EGFR provided by RHAMM^B^ and to understand whether endogenous levels of RHAMM^A^ activate EGFR signaling.

## Conclusion

We provide evidence that upregulation of RHAMM^B^ is a valuable prognostic marker for pancreatic cancer. We demonstrated that only the shorter isoform RHAMM^B^, but not RHAMM^A^ with 15 extra amino acids encoded by exon 4, is significantly upregulated in PNET and PDAC. RHAMM^B^, but not RHAMM^A^, promotes metastasis in spontaneous and experimental metastasis mouse models of PNET. EGFR signaling is required for RHAMM^B^-induced liver metastasis, but is not sufficient to promote PNET metastasis. *RHAMM*^*B*^, but not *RHAMM*^*A*^, is correlated with inferior survival in PDAC patients.

## Methods and Materials

### Bioinformatics and Statistical Analysis for *RHAMM* variants from RNA-Seq data

Total RNAs from 27 primary human PanNETs, 12 metastatic human PanNETs, and human BON1 cell line [8] were isolated with RNeasy Plus Universal Kits (Qiagen Cat no. 73404, Germantown, MD) according to the manufacturer’s protocol. Library preparation and RNA sequencing with paired-end 75 bp reads were performed according to protocols at the Genomics Resources Core Facility, Memorial Sloan Kettering Cancer Center, and Weill Cornell Medicine. A RNA-Seq dataset of 89 human pancreatic islets was obtained from publicly available database Gene Expression Omnibus (GEO accession GSE50398, http://www.ncbi.nlm.nih.gov/geo/query/acc.cgi?acc=GSE50398) [9-11], and was converted to fastq file from SRA format using SRA Toolkit (v 2.4.4). All samples analyzed were aligned to hg19 reference genome using STAR (v 2.4) aligner. Aligned samples were then quantified to obtain gene expression in terms of FPKM (Fragments per Kilobase of Transcripts per Million Reads) against a UCSC hg19 annotation Gene Transfer Format containing coordinates for all genes, including specific RHAMM isoforms of interest using CuffLinks (v 2.2.1). FPKM values were extracted from isoform specific quantification output obtained from CuffLinks for each sample, with focus on isoform A (NM_012484) and isoform B of *RHAMM* (NM_012485). Log-transformed FPKM values (log2 [FPKM+1]) were used for further analysis to compare the expression levels across tissue types (Normal, Primary and Liver metastasis) and *RHAMM* isoforms (*RHAMM*^*A*^ and *RHAMM*^*B*^) using two-way ANOVA, followed by pairwise comparisons with Tukey’s post-hoc test for multiple comparison adjustment. All analyses were performed in open-source data analysis software R (v 2.14.1) or SAS 9.4 (SAS Institute, Cary, NC).

Bioinformatics and statistical analyses were conducted on the publically available gene expression dataset from The Cancer Genome Atlas (TCGA; http://cancergenome.nih.gov/). TCGA normalized expression values were created by Illumina RNA-Seq version 2 RNA sequencing data (level 3). Downloaded data was analyzed using R or statistical software Prism (version 6.0f) for statistical computation and Kaplan-Meier survival analysis with log-rank test. Two-tailed Mann-Whitney U test was used to compare differences between two groups selected. P values < 0.05 were considered as statistically significant.

### Quantitative real-time reverse transcription PCR (RT-qPCR)

Messenger RNA (mRNA) was isolated from human PanNET specimens or cells grown on 6-cm or 10-cm plates using RNeasy mini kit (Qiagen) containing gDNA eliminator spin columns. The cDNA was generated using SuperScript III First-strand synthesis system with random hexamers (Invitrogen), and power SYBR green (Invitrogen)-based qPCR was performed with 3 internal control genes and the comparative C_T_ method (ΔΔC_T_).

The sequences of the primers used are *RHAMM*^*A*^ (forward: located within exon 3, 5’-TGACAAAGATACTACCT TGCCTGCT-3’, reverse: located at the junction of exon 3 and 4, 5’-TCATTCTTTTGAGATTCCTTTGATTC-3’); *RHAMM*^*B*^ (forward: located at the junction of exon 3 and 5, 5’-AAAGTTAAGTCTTCG GAATCAAAGATT-3’, reverse: located within exon 5, 5’-GCATTATTTGCA GAGAGAGATGT-3’), and internal control genes: human *HMBS* (forward: 5’-CCATCATCCT GGCAACAGCT-3’, reverse: 5’-GCATTCCTCAGGGTGCAGG-3’); human *EEF1A* (forward: 5’-CAATGTGGGCTTCAA TGTCAA-3’, reverse: 5’-CATAGCCGGCGCTTATTTG-3’); human *MRPL19* (forward: 5’-GGGATTTGCATTCAGAGATCAGG-3’, reverse: 5’-CTCCTGGACCCGAGGATTATAA-3’).

### Tissue preparation, immunohistochemistry, and scoring of protein expression

Retrospective and prospective review of PanNETs was performed using the pathology files and pancreatic cancer database at the authors’ institutions with Institutional Review Board (IRB) approval. Construction of TMA of human PanNETs was described previously [12]. Mouse tissues were fixed in 10% buffered formalin overnight at room temperature. Fixed tissues were processed and cut into 5 µm sections at Histoserv. Formalin-fixed/paraffin-embedded sections were deparaffinized and rehydrated by passage through a graded xylene/ethanol series before staining. Immunochemistry was examined by VECTASTAIN Elite ABC Kits (Vector Laboratories, Inc., Burlinggame, CA) following manufacturer’s instructions. The primary antibodies used were rabbit anti-RHAMM (1:1,000, Y800 [2], 1:100), rabbit anti-synaptophysin (1:100, Vector Laboratories, VP-S284 or 1:100, Lab Vision/neomarkers, Fremont, CA, RM-9111), and GFP (1:300, Invitrogen, Carlsbad, CA, A11122). RHAMM expression for each tumor was given a score of 0 if no staining was present, and a score of 1 if moderate to strong staining was present.

### Cloning of RCASBP and retroviral vectors

RCASBP is a replication-competent avian leucosis virus with a splice acceptor and the Bryan-RSV pol gene. RCASBP-*RHAMM*^*B*^ [2] and RCASBP-*EGFR\** [13] have been described. RCASBP-*RHAMM*^*A*^ was generated using QuikChange Lightning Site-Directed Mutagenesis Kit (Agilent Technology, Santa Clara, CA) to add exon 4 (forward primer: 5’-agttaagtcttcggaatcaaaggaatctcaaaagaatgataaa gatttgaagatattagagaaagagattcgtgttcttctacaggaac-3’, reverse primer: 5’-gttcctgtagaagaacacgaatctctttctctaatatcttcaaatctttatcattcttttgagattcctttgattccgaagacttaact-3’) from RCASBP-*RHAMM*^*B*^. The presence of exon 4 in RCASBP-*RHAMM*^*A*^ was confirmed by DNA sequencing.

### The shRNA knockdown

Hairpin sequences targeting *EGFR* are S7: CCAAGCCAAATGGCATATTTA and S9: GCTTTCGAGAACCTAGAAATA. Hairpin sequence targeting *LacZ* is TCGTATTACAACGTCGTGACT. Hairpin sequence targeting *RHAMM* is sh1: GCCAACTCAAATCGGAAGTAT (Sigma, Clone ID: NM_012484.2-2128s21c1). Lentiviruses harboring shRNA were generated using 293T cells as described previously (http://www.broadinstitute.org/rnai/public/resources/protocols).

### Cell culture and Western blot

Generation of N134 and BON1-TGL cell lines has been described [1, 14]. Human pancreatic neuroendocrine tumor cell line, BON1, was provided by Chris Harris [8, 12]. DF1, N134, and BON1-TGL were cultured in Dulbecco’s modified Eagle’s medium (DMEM) supplemented with 10% fetal bovine serum (FBS), 6 mM L-glutamine, and penicillin/streptomycin. The BON1-TGL cell line was transfected with pBABE-*RHAMM*^*B*^ with lipofectamine 2000 (Invitrogen) to generate BON1-TGL-RHAMM^B^ cell line. BON1-TGL cells overexpressing a control vector or *RHAMM*^*B*^ were cultured in DMEM supplemented with 0.5 µg/ml puromycin, 10% FBS, 6 mM L-glutamine, and penicillin/streptomycin. N134 cells overexpressing cDNAs were generated as described [15]. N134 cells overexpressing shRNAand BON1-TGL overexpressing shRNA were cultured in medium supplemented with 0.4 µg/ml puromycin and 0.5 µg/ml puromycin, respectively.

For Western blot analysis, cell extracts were loaded into mini-protean 4-15% pre-cast gels (Bio-Rad). Protein was immobilized onto nitrocellulose membrane, 0.45 µm pore size (Bio-Rad). Blots were blocked for one hour with 3% (weight/volume) bovine serum albumin (Fisher Scientific) and incubated overnight at 4° C with a rabbit monoclonal antibody to RHAMM [EPR4055] antibody at 1:1,000 dilution (Abcam, ab108339). The next day blots were washed 4 times for 10 minutes with TBST and incubated for one hour at room temperature with an anti-rabbit secondary antibody at 1:5,000 dilution. Bands were detected with enhanced chemiluminescence.

### Animal experiments

Generation of *RIP-Tag; RIP-tva* mice has been described [1] and detailed protocols for somatic delivery of the RCASBP viruses has been described [15]. Immunodeficient mice, NOD/scid-IL2Rgc knockout (NSG), were generated by the Jackson Laboratory. This study was carried out in strict accordance with the recommendations in the Guide for the Care and Use of Laboratory Animals of the National Institutes of Health. All mice were housed in accordance with institutional guidelines. All procedures involving mice were approved by the Institutional Animal Care and Use Committee of Weill Cornell Medicine. For the experimental metastasis assay, either 1 x 10^6^ N134 cells in 100 µL PBS were injected into the tail veins of NSG mice or 1 x 10^6^ BON1-TGL cells in 100 µL PBS were injected into the left ventricle of NSG mice. The orthotopic model of PanNET liver metastasis was performed as previously described [16] with the following modification: 0.5 x 10^6^ BON1-TGL cells in 50 µL DMEM containing 2% FBS were injected into the spleen. A standard formula for tumor volume was applied (volume [mm^3^] = 0.52 × width^2^ × length). Tumor burden is the sum of the tumor volume per mouse.

## Abbreviations

RHAMM: receptor for hyaluronic acid-mediated motility
PNET: pancreatic neuroendocrine tumor
TCGA: The Cancer Genome Atlas (TCGA)
HA: hyaluronic acid
FPKM: fragments per kilobase of transcripts per million reads
RT-qPCR: quantitative real-time reverse transcription PCR
PDAC: pancreatic ductal adenocarcinoma
DMEM: Dulbecco’s modified Eagle’s medium
FBS: fetal bovine serum.

## Declarations

### Ethics approval and consent to participate

All procedures involving mice were approved by the institutional animal care and use committee. Retrospective and prospective review of PNETs was performed using the pathology files and pancreatic cancer database at the authors’ institutions with IRB approval.

### Consent for publication

All authors agreed on the manuscript.

### Availability of data and material

The RNA-Seq and TCGA datasets that support the findings of this study are available in Gene Expression Omnibus (GEO accession GSE50398, http://www.ncbi.nlm.nih.gov/geo/query/acc.cgi?acc=GSE50398) [9-11], and http://cancergenome.nih.gov/cancersselected, and the other data are available form the authors upon reasonable request.

### Authors’ contributions

S.C., X.C., B.J.K., G.Z., S.P., T.H., and L.S. performed the experiments and analyzed the data. D.W. conducted the analysis on the TCGA dataset. D.W., G.Z., S.P., T.J. F.III, and K.R. edited the manuscript. L.H.T. contributed immunohistochemical interpretation. A.V., C.S.C., and O.E. contributed to bioinformatics analyses. Z.C. performed statistical analysis. L.H.T., K.R., and T.J. F.III contributed to samples. Y.-C.N.D. designed and performed the experiments, analyzed the data, and wrote the manuscript.

#### Acknowledgments

The authors thank Danny Huang for mouse database design; Harold Varmus, Yi Li, Jihye Paik, Todd R. Evans, David Foster, Bi-Sen Ding, Katie Politi, Jeffrey Yongchun Zhao, Romel Somwar, Selina Chen-Kiang, Stephanie Azzopardi, Samantha Li, Megan Wong, Joseph Na, and Robin Zhang for their valuable input and excellent assistance; Diane L. Reidy and David S. Klimstra for contributing the approval of the tumor materials.

---

**Table S1.** Impact of genes on lymph node and liver metastasis of PanNETs in *RIP-Tag; RIP-tva* mice.

| **RCASBP-** | **Age (week)** | **Lymph node metastasis** | **Liver metastasis** |
| --- | --- | --- | --- |
| *RHAMM*<sup>A</sup> | 16 | 0/8 mice (0%) | 0/8 mice (0%) |
| *RHAMM*<sup>B</sup> | 16 | 8/11 mice (73%) | 8/11 mice (73%) |
| *EGFR* * | 16 | 5/8 mice (62.5%) | 2/8 mice (25%) |

**Fig. S1.**
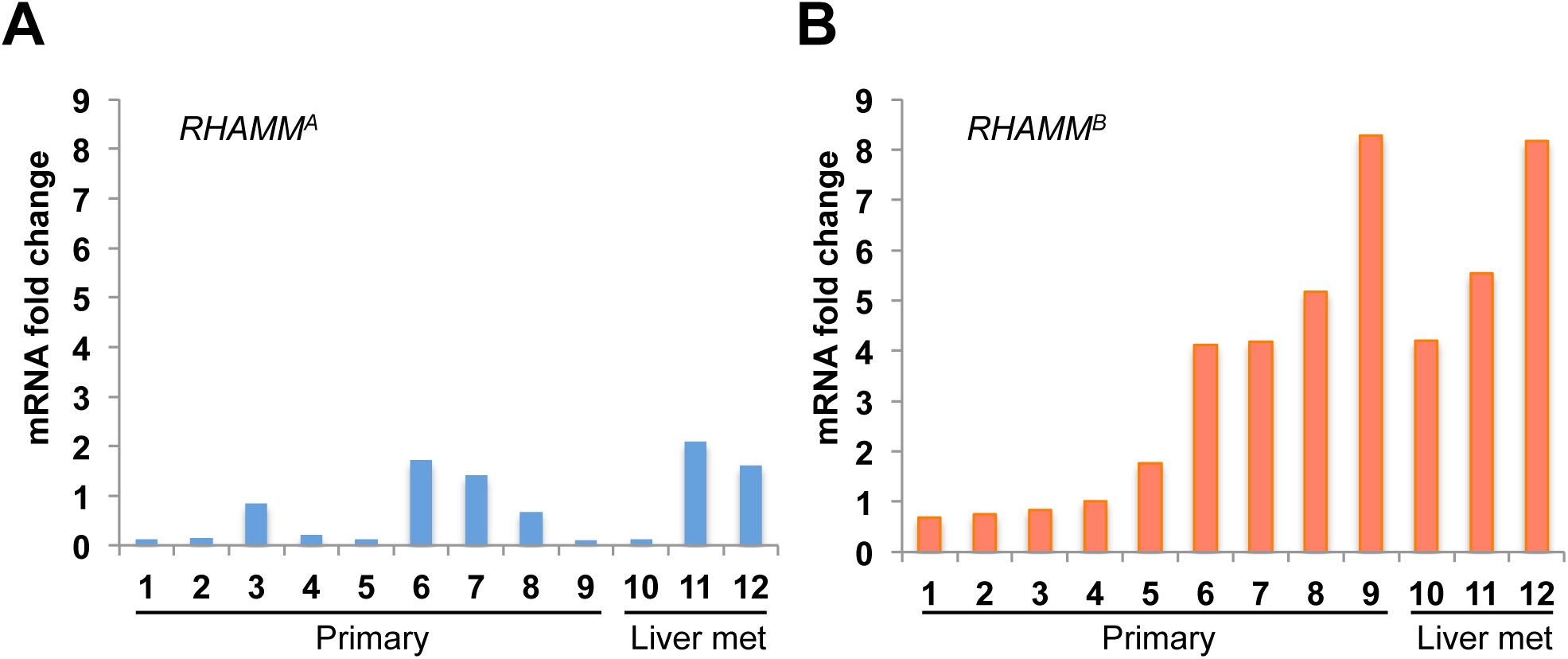
RT-qPCR analysis of *RHAMM*^*A*^ (A) and *RHAMM*^*B*^ (B) in 9 primary human PNETs and 3 metastatic PNETs from the livers.

**Fig. S2.**
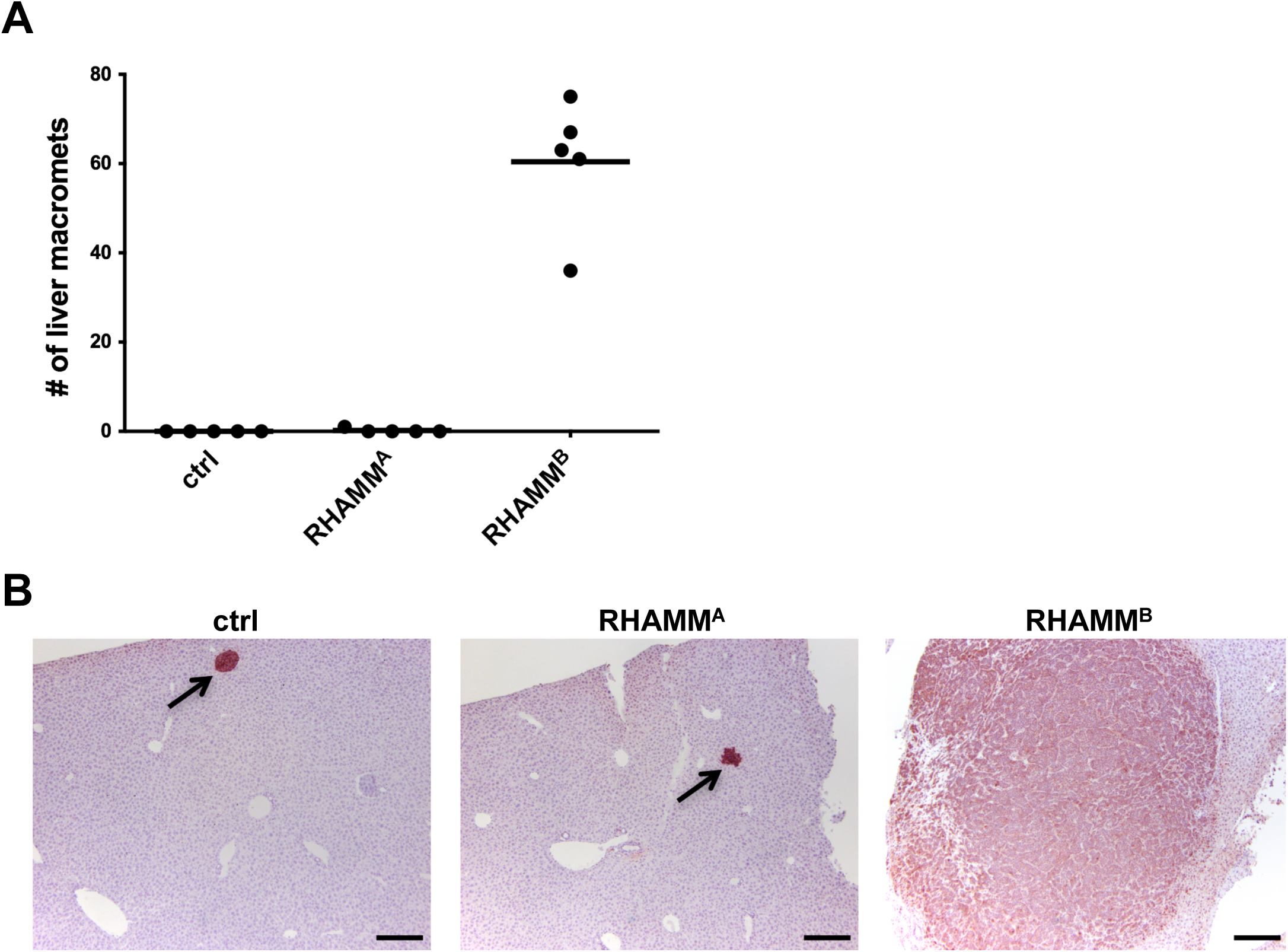
Unlike RHAMM^B^, RHAMM^A^ did not promote liver metastasis of mouse PNET N134 cell line in a tail vein experimental metastasis assay. (A) The number of liver macrometastases was recorded. (B) Immunohistochemical staining of synaptophysin in the liver sections to reveal the presence of metastatic PNETs. Arrows indicate micrometastases. Original magnification: 10X. Scale bar, 200 µm.

**Fig. S3.**
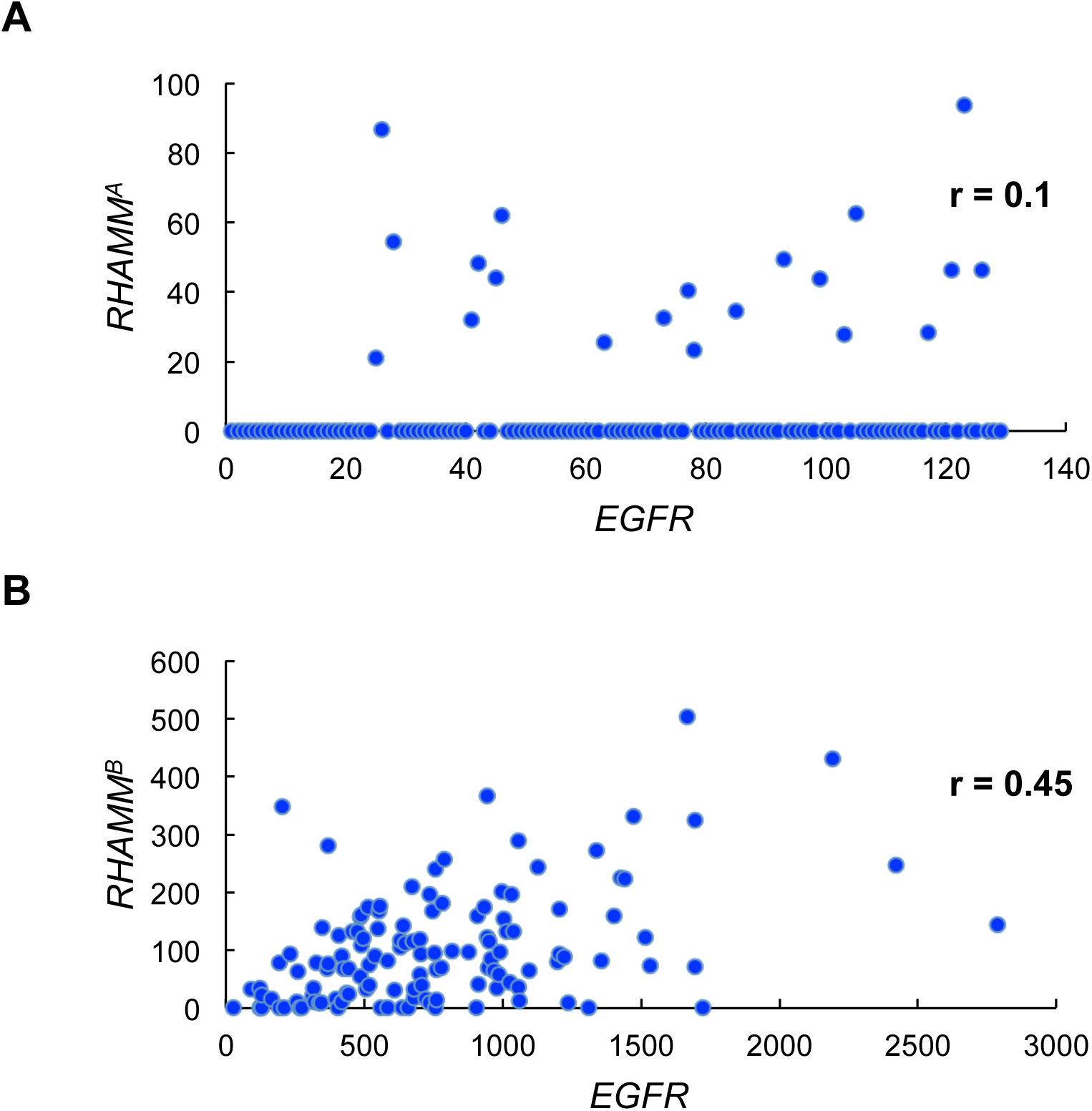
Correlation between *EGFR* expression and *RHAMM*^*A*^ (A) or *RHAMM*^*B*^ (B) expression in TCGA PDAC dataset (RNA-Seq V2).

